# Neural Basis of the Delayed Gratification

**DOI:** 10.1101/2021.03.10.434739

**Authors:** Zilong Gao, Hanqing Wang, Chen Lu, Sean Froudist-Walsh, Ming Chen, Xiao-jing Wang, Ji Hu, Wenzhi Sun

**Author notes:** Corresponding authors., &. These authors contributed equally to this work (Zilong Gao, Hanqing Wang).

## Abstract

Balancing instant gratification versus delayed, but better gratification is important for optimizing survival and reproductive success. Although psychologists and neuroscientists have long attempted to study delayed gratification through human psychological and brain activity monitoring, and animal research, little is known about its neural basis. We successfully trained mice to perform a waiting-and-water-reward delayed gratification task and used these animals in physiological recording and optical manipulation of neuronal activity during the task to explore its neural basis. Our results showed that the activity of DA neurons in ventral tegmental area (VTA) increases steadily during the waiting period. Optical activation vs. silencing of these neurons, respectively, extends or reduces the duration of waiting. To interpret this data, we developed a reinforcement learning (RL) model that reproduces our experimental observations. In this model, steady increases in DAergic activity signal the value of waiting and support the hypothesis that delayed gratification involves real-time deliberation.

**TEASER:** Sustained ramping dopaminergic activation helps individuals to resist impulsivity and wait for laerger but later return.

## INTRODUCTION

To optimize survival and reproductive success, animals need to balance instant gratification versus delayed, but better gratification. Repeated exposure to instant gratification may disrupt this balance, thereby increasing impulsive decisions. Such decisions contribute to numerous human disorders, such as addiction and obesity(*1, 2*). Delayed gratification is an important process that balances time delay with increased reward (*3*). It is influenced by strengths in patience, will-power, and self-control(*4*). Although psychologists and neuroscientists have long studied this important behavior through human psychological and brain activity assessments as well as rodent-based studies, little is known about its neurological basis.

During a well-controlled delayed gratification task, an individual must balance the benefits vs. risks of delay in receiving an available reward. Sustaining choice requires suppression of constant temptation by the expectation of enhanced reward in the future (*3, 5, 6*). Midbrain dopaminergic neurons are well known to play central roles in reward-related and goal-directed behaviors (*7-12*). Studies have revealed that DAergic activity signals proximity to distant rewards, either spatially or operationally (*7, 13, 14*), which has been postulated to sustain or motivate goal-directed behaviors while resisting distractions. DAergic neurons play important roles in time judgment (*15*) and cost-benefit calculations which are necessary for value-based decision making (*13, 16-18*).

We successfully trained mice to perform a waiting-and-water-reward delayed gratification task. Recording and manipulation of neuronal activities during this task allows us to explore the cellular regulation of delayed gratification. We found that the activity of VTA DAergic neurons ramped up consistently while mice were waiting in place for rewards. Transient activation of DAergic neurons extended and inhibition reduced the duration of the waiting period. Then we adopted reinforcement learning (RL) computational models to predict and explain our experimental observations.

## RESULTS

### Mice can learn to wait for greater rewards by delayed gratification task training

First, we trained mice to perform a one-arm foraging task (pre-training) in which delay did not result in increased reward (*19*). The period the mouse in the waiting zone was set as waiting duration and the time a mouse used in running from the waiting zone to fetch the water reward was as running duration (Fig. 1A, left panel). When a water-restricted mouse exited the waiting zone and licked the water port in the reward zone, it could receive a 10 µl of water drop regardless of the time spent in the waiting zone (Fig. 1A, right panel, black line). In a week of training, the average waiting and running durations both significantly decreased from Day 1 to Day 7 (p < 0.001, n = 7 mice, Figs. 1C-E, Movie S1). All mice learned the strategy of reducing durations of both waiting and running to maximize the reward rate (as ul of water per second in a trial, fig. S1A).

**Fig. 1.**
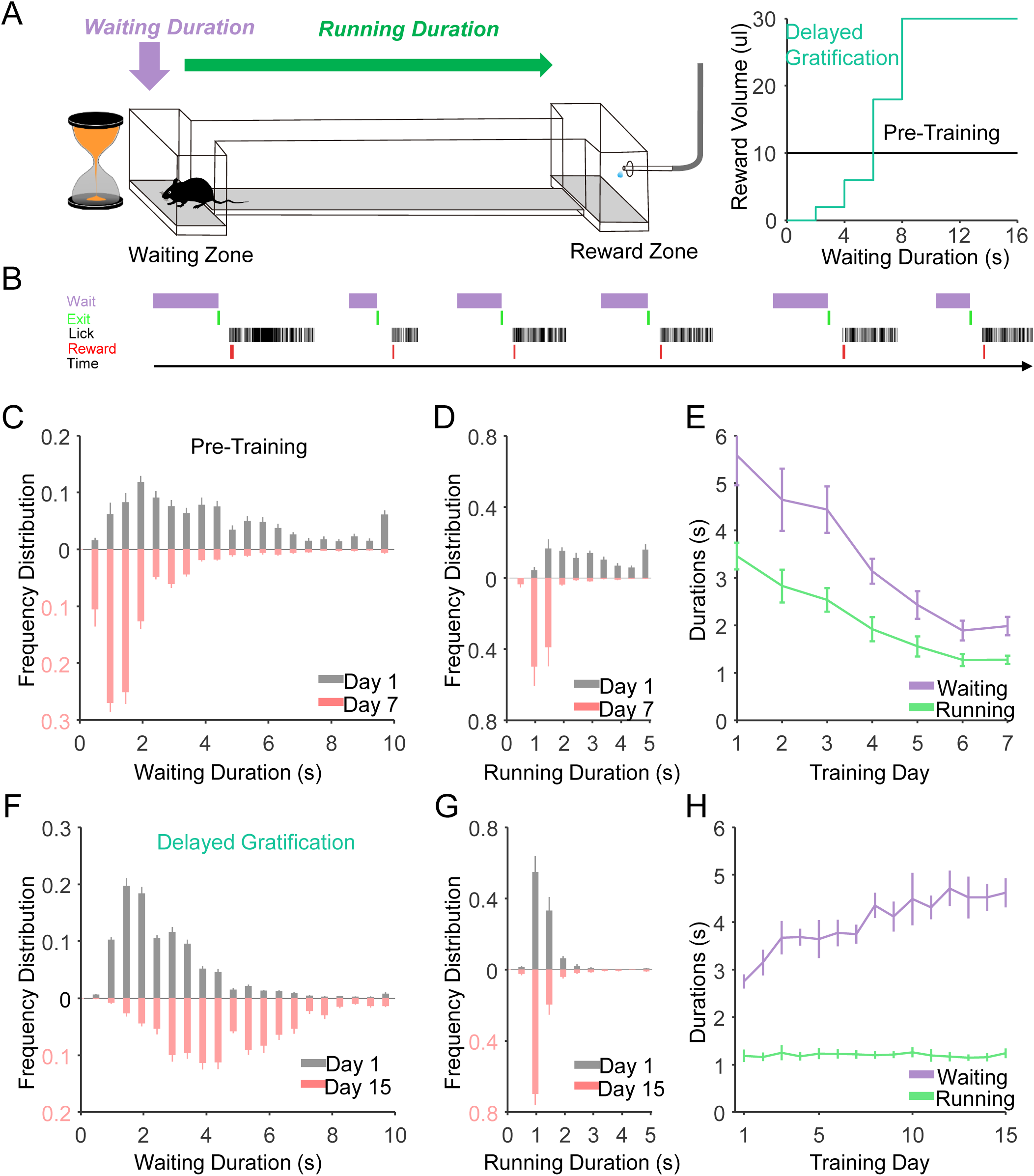
The behavioral performance of mice during a delayed gratification task learning. (**A**) Left panel, schematic of the delayed gratification task. Right panel, the relationship between reward volumes and waiting durations in the two behavioral tasks. (**B**) This plot presents the Transistor-Transistor Logic (TTL) signals for the chronological sequence of behavioral events in the tasks. (**C-E**). The waiting duration and running duration both decreased with the training process in the pre-training phase (Day1, Waiting: 5.58±0.63 sec; Running: 3.46±0.28 sec; p<0.001; Day 7, Waiting: 1.99±0.19 sec; Running: 1.28±0.09 sec, p<0.001, n=7 mice, Friedman test). (**F**) The distribution of waiting durations from the behavioral session on the last analyzed day (Day 15, light red), revealing significantly longer waiting durations compared with that from day 1 (D 1, gray, n=7 mice). (**G**) The distribution of running durations from D1 and D15 did not differ with training. (**H**) The plots show that the continuous training steadily increased the averaged waiting duration from 2.76 ± 0.15s on Day 1 to 4.62 ± 0.30 s on Day 15 (p < 0.001, n = 7 mice, Friedman test), whereas the training did not change the average running duration from 1.19±0.13s on Day 1 to 1.24 ± 0.10 s on Day 15 (p = 0.97, n = 7 mice, Friedman test). All error bars represent the s.e.m..

Next, we trained the same mice using a delayed gratification paradigm, where the size of the reward increased quadratically with time spent in the waiting zone (Fig. 1A, right panel, green line). Over the next three weeks, this resulted in shifting of the distributions of waiting duration towards longer time durations. The averaged waiting period significantly increased from Day 1 to Day 15 (p<0.001, Figs. 1F, H, and Movie S1), whereas the duration of running did not decrease beyond that observed initially (p = 0.97, n = 7 mice, Fig.s 1G, H). The reward rate increased steadily, indicating that the mice were learning to successfully delay gratification (fig. S1B).

### The activity of VTA DAergic neurons increases steadily during the waiting period

To monitor the activity of VTA DAergic neurons during the delayed gratification task, we employed fiber photometry to record the calcium signals in VTA DAergic neurons in freely moving mice for as long as one month (Fig. 2A-C, optical fiber placement illustrated in fig S2). On the first day of pre-task training, the calcium signal rose rapidly on reward and quickly reached a peak. A few days of training dramatically reshaped the response pattern. Once the mice re-entered the waiting zone, the activities of DAergic neurons started to rise and reached the highest level when the animal received a reward (fig. 3SA).

**Fig. 2.**
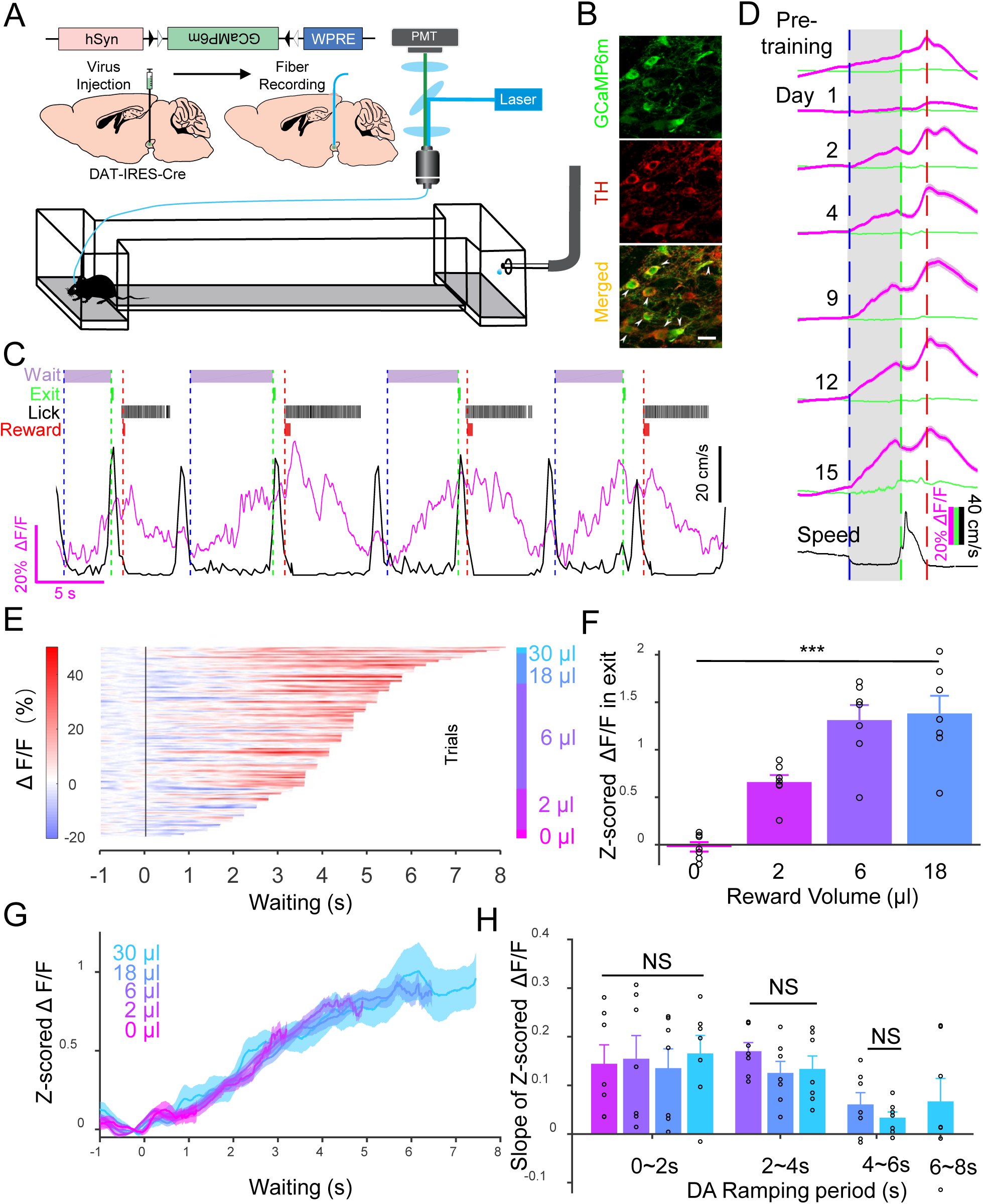
VTA DAergic activity ramps up consistently while the mice are waiting for the reward. (**A**) Schematic of stereotaxic virus injection procedures. (**B**) Confocal images illustrating GCaMP6m (green) expression in VTA TH^+^ neurons (red). Scale bar: 20 μm. (**C**) An example of a live-recording trace (magenta line) of Ca^2+^ signal in VTA ^DA^ neurons and running speed (black line) when a Dat-Cre: GCaMP6m mouse was performing the delayed gratification task. Delayed gratification task events over time (top): the dashed vertical lines indicated waiting onset (blue), waiting termination (green), and reward onset (red). (**D**) The scaled Ca^2+^ signals curves (magenta) and GFP signals (green) curves of VTA^DA^ neurons from the last day in pre-training and day 1 to day 15 in the delayed gratification task training (black line, speed). (**E**) Sorted Ramping Ca^2+^ signal data from one mouse on the last day (D15) of the delayed gratification task training (150 trials). The signal traces were aligned to waiting onset, sorted in waiting duration length, and separated into five groups of the reward outcomes (0, 2, 6, 18, and 30μl). **f**. Z-scored ΔF/F values at 0.5s before exit were significantly different while the reward volumes were different (F=24.67, p<0.01, n=7, one-way ANOVA). (**G**) Averaged Ca^2+^ signal curves with different outcomes from **Fig. E**. Slopes of Ca^2+^ signals for every outcome, showing that there were no differences in all DAergic ramping periods throughout the last week of training (0∼2s, F=0.10, p=0.96; 2∼4s, F=1.03, p=0.38; 4∼6s, F=1.00, p=0.34, n=7, one-way ANOVA). All error bars represent the s.e.m.. For (**D**) and (**G**), the shaded region represents s.e.m..

**Fig. 3.**
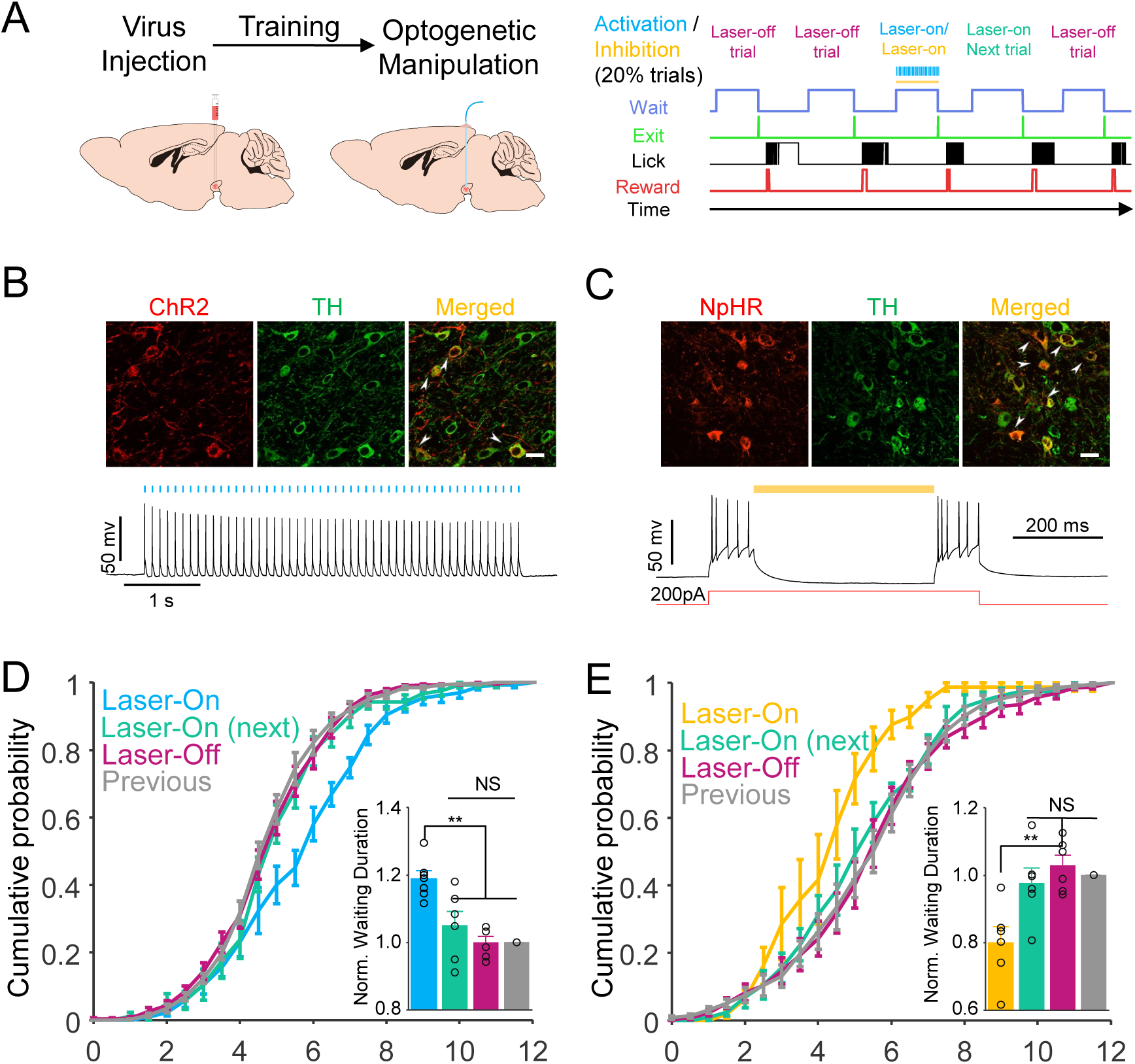
Optogenetic manipulation of VTA DAergic activity altered the waiting durations. (**A**) Left panels: schematic of stereotaxic virus injection and surgical procedure. Right panels: the behavioral events and optogenetic manipulation protocol. (**B**) Top panels: confocal image showing ChR2-mCherry (red) expression in VTA TH+ neurons (green). Bottom panels: whole-cell recording of VTA TH+ neurons in brain slice showing action potentials evoked by 10-Hz 473 nm laser flash sequences (50 flashes, 20ms interval). (**C**) Top panels: confocal image showing eNpHR3.0-mCherry (red) expression in VTA TH+ neurons (green). Bottom panels: whole-cell recording of VTA TH+ in brain slice showing that action potentials evoked by 200pA current injection were inhibited by continued 589nm laser. Scale bar: 20 μm. (**D**) Cumulative probabilities of waiting durations. The waiting durations of optogenetically activated trials were significantly increased (blue, F=12.93, p=0.002, n=6 mice, one-way ANOVA) than that of the previous day’s trials (gray); note that the waiting duration of unstimulated trials (red) did not differ from that of the previous day’s trials (magenta, p=0.96), or the next trials following photoactivation (green, p=0.63). Insert: a bar graph of the normalized waiting durations from lasing stimulation (blue), the previous day’s trials (gray), photoactivated trials (blue, 1.19±0.03), unstimulated trials (red, 1.00±0.02), and the next trials following the photoactivation (green, 1.05±0.04). (**E**) The same experimental configuration as in (**D**), but VAT TH^+^ neurons were optogenetically inhibited by a yellow laser. Optogenetic inhibition decreased the waiting duration (yellow, 0.80±0.05, F = 7.76, p=0.008, n=6 mice, one-way ANOVA), whereas there was no difference between uninhibited trials (red, 1.03±0.03, p=0.80), the trials following the photoinhibition (green, 0.98±0.04, p=0.80), and the previous day’s trials (gray). All error bars represent the s.e.m..

We next analyzed the activity of these same neurons in the mice as they learned the delayed gratification task. The recording traces showed that training gradually reshaped the pattern and time course of activity (Fig. 2D). The activity started to ramp up once the mice entered the waiting zone, and then reached its highest level when animals exited. To investigate carefully the dynamical properties of the ramping activity during waiting, we sorted the calcium signals from one training day of one mouse by their length of waiting durations and plotted them with a heat map (Fig. 2E). We divided trials according to the trial outcome (reward volume) and calculated the calcium signals while the mouse exited waiting zone with different reward volumes. Our results showed that the Z-scored calcium signals at 0.5 sec before exit were significantly different while the reward volumes were different (Fig. 2F), but the mean signal curves raised along a similar trajectory regardless of trial outcome (Fig. 2G). Then, we calculated the slopes of signal curves with different outcomes over 4 time windows (0∼2, 2∼4, 4∼6, 6∼8 s) by linear regression analysis. The slopes during the same time window had no significant differences between reward groups (Fig. 2H). We pooled and plotted the slopes of different waiting periods together and found the activity curves kept rising steadily and almost saturated after 6 secs from the time the mice entered the waiting zone. Besides, the ramping DAergic activity became less variable along with delayed gratification task training in our experimental data (figs. S4A-D). All these results indicated the VTA DAergic neurons consistently ramp up their activity during waiting in as animals are trained in the delayed gratification task.

### Optogenetic manipulation of VTA DAergic activity altered the waiting durations in delayed gratification task

To determine whether VTA DAergic activity controls performance in the delayed gratification task, we manipulated VTA DAergic neurons temporally within 20% pseudo-randomly chosen trials utilizing optogenetic tools while the mice were waiting during the delayed gratification task (Figs. 3A-C). Activating the VTA DAergic neurons shifted the cumulative probability distribution to statistically significant longer waiting duration (Fig. 3D, blue), while inhibiting these same neurons shifted this distribution to significantly shorter periods(Fig. 3E, yellow). The impacts on the cumulative probability duration distributions were only observed in the Laser-ON trials. In contrast, the Laser-OFF trials, including the next trials after the Laser-ON as a single group, were not significantly different from the trials from the previous day (Figs. 3D-E). The optical manipulations didn’t influence the running durations in ChR2 or eNpHR 3.0 expressing mice (figs. S5A-B), nor did it change the waiting duration distribution of mice that expressed mCherry in DAergic neuron in delayed gratification task (figs. S6A-B). To rule out the possibility of optogenetic manipulation-induced memory, we performed a random place preference test with the same stimulation dosage. Nither activating nor inhibiting VTA DAergic neurons significantly changed the transient waiting duration and pattern in the location in which the laser was activated in all tested mice (figs. S5C-F) as well as the mCherry expressing controls (figs. S6C-F).

### A reinforcement learning (RL) model suggests that ramping VTA DAergic activity signals the value of waiting for delayed gratification

How does a mouse manage to wait longer for a larger reward vs. smaller but more immediate reward options? We propose two models to explain behavioral scenarios to exemplify possible strategies a mouse may implement to achieve extended waiting performances: setting a goal of expected waiting duration before initiation of waiting or continuously deliberating during the waiting period. According to the first hypothesis, we modeled a RL agent that keeps timing until the preset moment has passed(Fig. 4A, *Decision Ahead*); In the second, we modeled a second RL agent that continuously balances the values of waiting versus leaving to control the decision on waiting versus leaving for the reward. Practically, we used a version of the state-action-reward-state-action (SARSA) algorithm with a series of stares (*20, 21*)(Fig. 4A, *Continuous Deliberation*, see methods). The behaviors of both models were able to replicate the behavioral performance we observed in animal experiments (Figs. 4B-C). There was no significant difference between the Kullback-Leibler (KL) divergence we chose to quantitatively assess the divergence from the waiting distribution of simulated behavior to that of the animals for both RL models. We couldn’t determine which model is better based on behavioral performance alone, given that both models well reproduced the behavioral data (Fig. 4D).

**Fig. 4.**
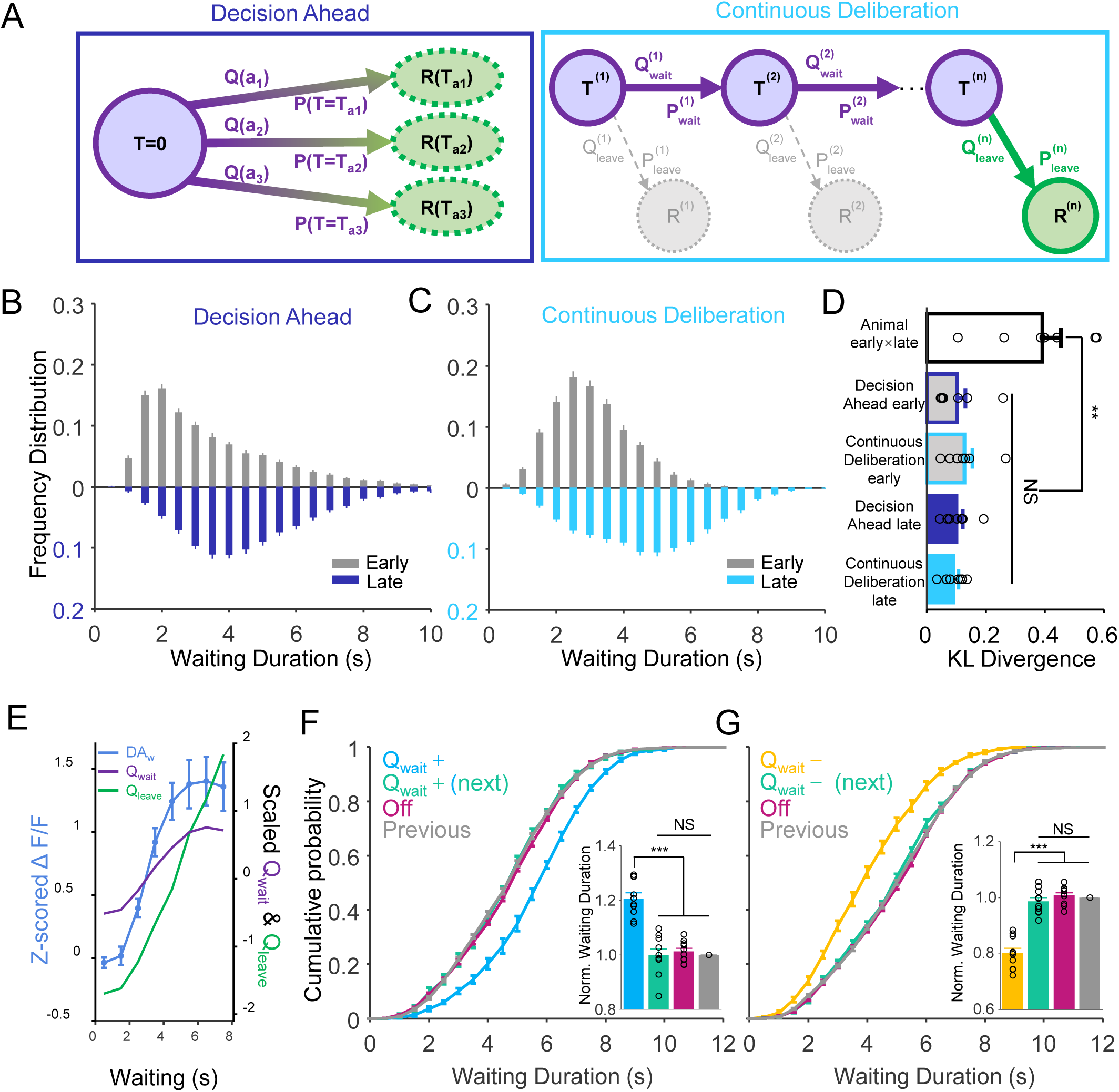
Behavioral performances and ramping VTA DAergic activity are explained by RL model. (**A**) Two reinforcement learning computational models, *Decision Ahead* in the dark blue box and *Continuous Deliberation* in the light blue box, simulating the decision processes and variables under the delayed gratification task. In *Decision Ahead*: Q(a_n_) is the value for action a(n) and P(T_an_) is the probability of action a(n); Q(a_n_) was used to compute the probability of action a(n). In *Continuous Deliberation*: probability of waiting,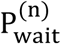; waiting action value, 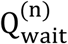; probability of leaving, 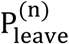; leaving action value, 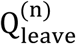; R^(n)^, received reward; 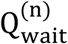 and 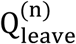 were used to compute the probability of waiting or leaving. (**B-C**) The distributions of waiting durations from the early session and late session simulated in *Decision Ahead* model (**B**) and *Continuous Deliberation* model (**C**) both displayed a similar distribution with our experiment data (**Fig. 1F)**. (**D**) The distributions of behavioral performances between early training days and late training days from experiment data were very different, in which the Kullback-Leibler (KL) divergence was big enough (0.39 ± 0.06). The KL divergences between the distributions of simulated behavioral performances from both models in early or late session and experiment data of every mouse cross whole training sessions were significant small (p = 0.005, n = 7 mice, Friedman test), in which there was no difference (p > 0.99) between *Decision Ahead* RL model and *Continuous Deliberation* RL model in late and early session. Data are represented as mean ± SEM. (**E**) Plots of Z-scored ΔF/F values (DA_w_, light blue) at 0.5s before the waiting ended, the scaled values of waiting (Q_wait_, light purple), and the value of leaving (Q_leave_, green) from *Continuous Deliberation* model in last training session. The Q_wait_ and Q_leave_ both predicted the experimental observation well (Q_wait_: r = 0.99, p<0.001; Q_leave_: r = 0.91, p = 0.002, Pearson correlation). (**F-G**) Computational reinforcement learning model (continuous deliberation)-simulated data, dependent on manipulating the value of waiting (Q_wait_) in delayed gratification task. As with the experimental data in **Fig. 3D-E**, the model-simulated data also shows that increased Q_wait_ only increases the waiting durations of Q_wait_ increased trials ((**F**), p<0.001, Friedman test, n=10) whereas decreased Q_wait_ can decrease the waiting durations of Q_wait_ decreased trials ((**G**), p<0.001, Friedman test, n=10). The unstimulated trials including the next trials after Q_wait_ manipulation had no difference with the last round regular running ((**F-G**), p>0.999, Friedman test, n=10). All error bars represent the s.e.m..

What does the ramping DAergic activity mean in the delayed gratification task? We tried to explain it with our RL model. In the *Decision Ahead* model, the agent keeps timing until the preset moment has passed, which suggests the ramping DAergic activity may relate to timing in delayed gratification task. Some studies have proposed that the ramping activity is consistent with a role in the classical model of timing with the movement initiated when the ramping activity reaches a nearly fixed threshold value, following an adjusted slope of ramping activity(*22-25*). In contrast, our results showed that the DAergic activity ramped up to different values with similar trajectories on a nearly constant slope (Figs. 2F-H). This suggests that VTA DAergic neurons may not implement a decision variable for the *Decision Ahead* scenario. In the *Continuous Deliberation* RL model, we compared the curves of the value of waiting and leaving with the ramping DAergic activity and found the behavioral performance of both animals and model agents reached the asymptote. The values of waiting (Fig. 4E, light purple) and the leaving (Fig. 4E, green) each correlated positively with the ramp of DAergic activity during waiting (Fig. 4E, green, blue, Z-scored ΔF/F, 0.5 sec before exit from the last week of training). This detailed analysis suggested that the *Continuous Deliberation* RL model agreed with previous studies (*13, 26-28*) and that ramping DAergic activity signals the value of actions, either waiting or leaving, in the delayed gratification task.

In the *Decision Ahead* RL model, if the agent keeps timing during waiting through ramping DAergic activity to encode the elapse of time (*29-31*), extra VTA DAergic activation should represent a longer time thus lead to an earlier stop of waiting. This deduction is contrary to our optogenetics result, namely DAergic activation led to a longer waiting (Fig. 3D). Instead, we reproduced the optogenetic manipulations in the *Continuous Deliberation* RL model by either increasing or decreasing the value of waiting (Q_wait_) in pseudo-random 20% trials. The increase or decrease in waiting durations only occurred in the Q_wait_-manipulated trials, whereas the remaining trials, including the next trials after value manipulation, had no significant difference with control (Figs. 4F-G). Importantly, manipulating the value of leaving (Q_leave_) in pseudo-random 20% trials induced the opposite results (figs. S8A-B) compared with experimental data of optogenetic manipulation. Our experimental data and *Continuous Deliberation* RL model together indicated that the ramping VTA DAergic activity profoundly influenced the waiting behavioral performance in the delayed gratification task, which suggested ramping DAergic activity signal the value of waiting, rather than the value of leaving. Our analysis conceptually revealed that the delayed gratification involved real-time deliberation.

### VTA DAergic activity during waiting predicts the behavioral performance in the delayed gratification task

Our optogenetic manipulation experiments and *Continuous Deliberation* RL model indicated that VTA DAergic activity during waiting influenced the waiting durations while the mouse was performing the delayed gratification task (Figs. 3D-E and Figs. 4F-G). Although the activity of VTA DAergic neurons ramped up consistently during waiting (Figs. 2G-H), they still fluctuated to a certain extent moment by moment. Therefore, we are next to determine whether this fluctuation influences the waiting behavior in the delayed gratification task. A strong prediction given by the *Continuous Deliberation* model is that, if DAergic activity signals the value of waiting at each specific moment, the more likely the agent will keep waiting in the next “time bin”, but not in the later ones (fig. S8E-F, the value of waiting is only positively correlated with the behavior of next time bin), which agrees with the Markov property(*32*). We thus aimed to test the relationship between the amplitude of momentary VTA DAergic signal and the behavior (i.e., waiting or leaving) within each time bin to determine how the momentary DAergic activity (the calcium signal amplitude in 0∼1 sec, 1∼2 sec, 2∼3 sec, or 3∼4 sec after waiting onset, shown as each cluster of bars in Fig. 5B) affects the waiting performance in the subsequent periods (behavior within 1∼2 sec, 2∼3 sec, 3∼4 sec, and 4∼5 sec for DA in 0∼1 sec, behavior within 2∼3 sec, 3∼4 sec and 4∼5 sec for DA in 1∼2 sec, and so on, Fig. 5A). To integrate data from multiple sessions as well as multiple animals, we took the advantage of the linear mixed model analysis (LMM, or linear mixed-effects, LME, see method) (*33-35*). The regression coefficients between momentary DAergic activities and momentary waiting (1 for waiting and 0 for not) were significantly positive between adjacent DAergic and behavioral periods (Fig. 5B, 1sec for the bars on the most left of each cluster/adjacent DAergic-behavior). The pair of momentary DAergic activity in 3∼4 sec and waiting in 4∼5 sec didn’t show a significant correlation (p = 0.61, n = 7), which may possibly result from insufficient data for those long trials. This result indicates that the waiting decision of the current moment is only influenced by the most recent DAergic signal but not by DAergic signal further in the past, which suggests that deliberation for waiting in delayed gratification may be a Markov process as we formalized in the *Continuous Deliberation* RL model(*32*).

**Fig. 5.**
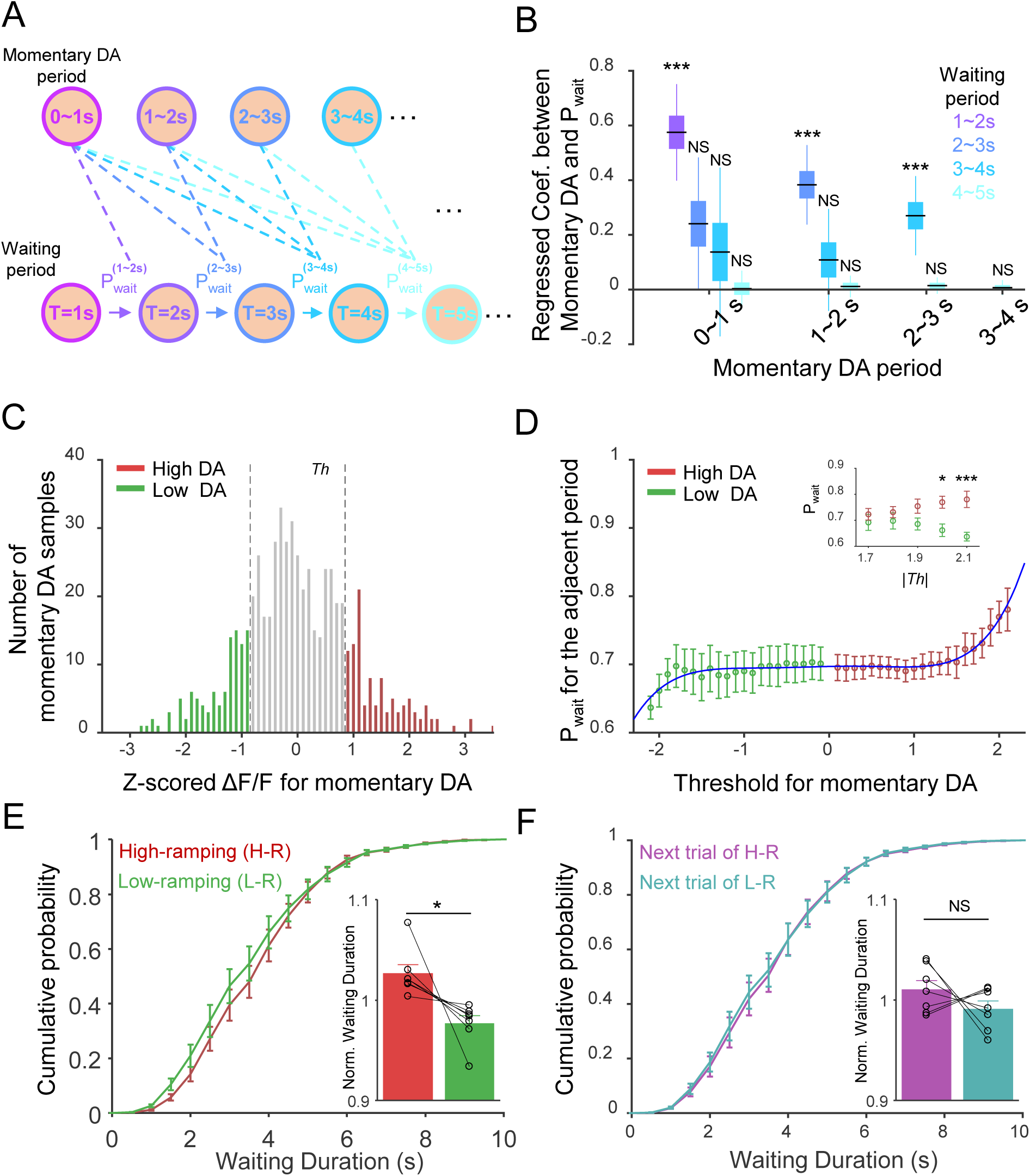
VTA DAergic activity during waiting predicts the behavioral performance in the delayed gratification task. (**A**) Schematic of waiting probability (P_wait_) in waiting periods after momentary DAergic periods from the experimental data. For momentary DAergic activity in each period, the mouse has P_wait_, which is calculated by the waiting durations and trial number, in any waiting periods after the momentary DAergic period. (**B**) Relationship between momentary VTA DAergic activity (Ca^2+^ signals) and its waiting probability. For each momentary DAergic period, its DAergic activity is only highly correlated (p<0.001, n = 7 mouse, black lines, regressed coefficient median; boxes, 50% confidence interval; whisker, 95% confidence interval) with P_wait_ in the adjacent waiting period (the left bar of each cluster). (**C**) The distribution of Z-scored mean ΔF/F of momentary DAergic periods. Three colors illustrate high dopamine activity (High DA: red, greater than the threshold value, gray dash line, while the threshold value is positive) trial numbers, low dopamine activity (Low DA: green, less than the threshold value, while the threshold value is negative) trial numbers, and all other (gray) dopamine activity trial numbers. (**D**) The waiting probability of High DA (red) and Low DA (green) activity trials for the adjacent period after the momentary DA periods. The P_w_ of High DA and Low DA activity trials fit well with a fifth-degree polynomial function (R^2^=0.93, −2.1≤ *threshold*≤2.1). While the absolute values of the threshold are big enough (|*Th*|≥1.7), the P_w_ of the High DA activity trails is significantly (p=0.04, F(1,12)=5.483, Two-way ANOVA) higher than the P_wait_ of the Low DA-ramping activity trials in adjacent waiting periods (|*threshold*|**=**2.0, p=0.02; |*threshold*|**=**2.1, p<0.001, Sidak’s multiple comparisons test, n=7). (**E-F**) Cumulative probabilities of waiting durations for the high DA-ramping trials (**e**, H-R, red), the lower-DA-ramping trials (**E**, L-R, green) and their “next-in-series” trials (**F**). Bar graph showing that the normalized waiting durations (1.03±0.01) of the higher-DA-ramping trials are significantly longer than that of the lower-DA-ramping trials ((**E**), 0.98±0.01, p=0.024, n=7, paired Student’s t-test), but have no difference between their “next-in-series” trials ((**F**), next trial of H-R, 1.01±0.01; next trial of L-R,0.99±0.01; p=0.290, n=7, paired Student’s t-test). All error bars represent the s.e.m..

In the *Continuous Deliberation* RL model, the probability of waiting (P_w_) positively correlates with the value of waiting (Q_wait_). To explore the impact of DAergic activity on the probability of waiting in our experimental data, we binned DAergic activity of every trial and normalized data points (V_DA_) in each momentary DAergic period (1 sec started from 0 to 9 s). Then, we divided the trials into two groups by setting a series of arbitrary thresholds (red, High DAergic activity, V_DA-Z_ ≥ *Th*; green, Low DAergic activity, V_DA-Z_ ≤ *-Th, Th* was the threshold for the analysis of high/low DAergic activity) from these trials (Fig. 5C, *Th* was set to 0.9). By analyzing the probability of waiting of low and high DAergic activity at different thresholds for the adjacent waiting period, we found that the probability of waiting increased rapidly as the absolute value of threshold was set larger and larger. The probability of waiting was significantly different between the high and low dopamine trials when the absolute value of *Th* (|*Th*|) was greater than 2.0 (Fig. 5D).

Finally, we investigated the influence of fluctuations of intrinsic VTA DAergic activity on the waiting performance of mice in the delayed gratification task. There were trials whose DAergic activity in the whole waiting duration was significantly higher (red, high-ramping) or lower (green, low-ramping) than the mean DAergic activity (see Methods). Then we separated two groups of trials and found that the cumulative distribution of waiting durations of high-ramping trials shifted to the right with significantly higher normalized waiting durations compared with the normalized waiting durations of low-ramping trials (Fig. 4E), but there was no difference between the normalized waiting durations for the next-in series trials of high-ramping and low-ramping trials (Fig. 4F). These results accorded with our optogenetic manipulation experiment (Figs. 3D-E) that optogenetically manipulated VTA DAergic activity transiently influences the behavioral performance of waiting in delayed gratification task.

## DISCUSSION

Here we reported a novel behavioral task to train the mice to learn a foraging task with a delayed gratification paradigm. Mice learned to wait for bigger rewards with the increase of waiting durations (Figs. 1F-H). Moreover, the calcium signal of VTA DAergic neurons ramped up consistently when the mouse waited in place before taking action to fetch an expected reward (Figs. 2G-H). Further data analysis showed that the ramping VTA DA activity indeed influenced the behavioral performance of waiting (Figs. 5B-E), which was confirmed with bi-directional optogenetic manipulations of VTA DAergic activity (Figs. 3D-E). At last, a RL model well predicted our experimental observations and consolidated the conclusion that the ramping VTA dopaminergic activity signaled the value of waiting in the delayed gratification task, which involves real-time deliberation (Figs. 4B-G)..

DA release in NAc was previously conjectured for sustaining or motivating the goal-directed behavior as well as resisting distractions (*13, 14*). Here, we explicitly implemented continuous ‘distractions’ or less-optimal options along the delayed gratification process, in which, to achieve better performance, the mice need to sustain waiting as well as prevent/control impulsivity (*3, 6, 36, 37*). We found remarkable and sustaining DAergic activation when mice managed to wait longer, and further demonstrated a causal link between DAergic activation and the increase in transient waiting probability. Furthermore, we found DAergic activity ramps up in a consistent manner during waiting, mimicking the value of waiting along with a series of states in our *Continuous Deliberation* RL model, both of which are presumably resulted from and contributed to resisting an increasing magnitude of distraction in our task. Intriguingly, the momentary DAergic activity was found positively correlated to the momentary waiting probability, which also suggested DAergic activity may be involved in the continuous deliberation process. Therefore, we not only for the first time to our knowledge demonstrated the behavioral significance of DAergic activity in delayed gratification, but also depicted a “Continuous Deliberation” framework where DAergic activity may participate and help achieve more flexible and sophisticated performance.

Numerous works use Pavlovian conditioning in studying DA activity(*10, 12, 38-40*). Some studies paired the reward with a cue (or cues), in which animals don’t need effortful work to obtain rewards. It is well known this kind of DA activity signals the RPE via phasic firing. In the studies using operant conditioning or goal-directed behavior, the animals have to perform actions and need effortful work to obtain outcomes, and a ramping DA activity was reported to emerge while the animals were approaching the reward (*13, 14, 41, 42*). The ramping activity is suggested to signal the value of work (*13*) or distant rewards (*14*), but key evidence is lacking because the change of sensory input flow remarkably alters the DA activity over time. Under such mutual influence, it is impossible to identify RPEs or the value of work from external cues. The RPE model of ramping activity assumes that the value increases exponentially (or at least in a convex curve) as the reward is approached. Under this model, sensory feedback is suggested to result in the RPE signal to ramp (*41, 43, 44*), while a lack of sensory feedback is predicted to make a flat RPE signal. In contrast, the ramping DAergic activity is well isolated from the external sensory inputs when performing a delayed gratification task in our model. The mice continuously deliberate the current state and future rewards without any external sensory inputs during waiting in place. We still observed the calcium signal of VTA DAergic neurons ramped up in a stable dynamic. This ramp may indicate an escalating value for the closer reward in temporal and represent the ‘willpower’ of waiting.

Midbrain DAergic neurons play an important role in reinforcement learning(*9, 11, 12, 45, 46*), where activation of DAergic neurons usually produces a reinforcement effect on associated action, stimulus, or place. But in our delayed gratification task, optogenetic manipulation of DAergic activity substantially influenced the ongoing behavior on the current trial without visible reinforcement effect on later trials. Notably, this optogenetic manipulation was not sufficient to induce a reinforcement effect in the random place performance test. These results revealed the distinct and potent instantaneous effect of DAergic activity during delayed gratification. The observations and analysis in our experiments integrate more reliable evidence for the value coding in VTA DAergic neurons and significantly update the understanding of the coding mechanisms and fundamental functions of the DAergic system in delayed gratification. Our design of the delayed gratification task recapitulates the realistic situation where distractions and less-valuable choices lie in the way of pursuing a larger but later benefit.

The deficit of resisting distractions (temptations), which disrupt the balance between constant reward and delayed reward, is closely related to a variety of disorders like obesity, gambling, or addiction(*1, 47*). The ramping VTA DAergic activity accords with the model about NOW vs LATER decisions that tonic/stable DAergic signal have a strong influence on dlPFC and favor LATER rewards(*2*). We proposed that the sustained VTA DAergic activity during the delayed period could serve as a conservative neural basis for the power to resist the ubiquitous distractions (temptations) and improve reward rate or goal pursuit in the long run.

## MATERIALS AND METHODS

### Mice

Animal care and use strictly follow institutional guidelines and governmental regulations. All experimental procedures were approved by the IACUC at the Chinese Institute for Brain Research (Beijing) and ShanghaiTech University. Adult (8-10 weeks) DAT-IRES-Cre knock-in mice (Jax stock# 006660) were trained and recorded. Mice were housed under a reversed 12/12 day/night cycle at 22–25°C with free access to ad libitum rodent food.

### Stereotaxic viral injection and optical fiber implantation

After deep anesthesia with isoflurane in oxygen, mice were placed on the stereoscopic positioning instrument. Anesthesia remains constant at 1∼1.5% isoflurane supplied per anesthesia nosepiece. The eyes were coated with aureomycin eye cream. The scalp was cut open, and the fascia on the skull was removed with 3% hydrogen peroxide in saline. The Bregma and Lambda points are used to level the mouse head. A small window of 300∼500µm in diameter was drilled just above VTA (AP: −3.10 mm, ML: ±1.15mm, and DV: −4.20 mm) for viral injection and fiber implantation. 300 nl of AAV2/9-hSyn-DIO-GCamp6m (10^12) solution was slowly injected at 30nl /min unilateral for fiber photometry recording. 300 nl either AAV2/9-EF1a-DIO-hChR2(H134R)-mCherry (10^12) or AAV2/9-EF1a-DIO-eNpHR3.0-mCherry (10^12) was injected bilaterally for optogenetic experiments. The injection glass pipette was tilted with an angle of 8° laterally to avoid the central sinus. After injection, the glass pipette was kept in place for 10 min and then slowly withdraw. An optical fiber (200 μm O.D., 0.37 NA; Anilab) hold in a ceramic ferrule was slowly inserted into the brain tissue with the tip slightly above the viral injection sites. The fiber was sealed to the skull with dental cement. Mice were transferred on a warm blanket for recovery, then housed individually in a new home until all experiments were done.

### Behavioral tasks

One week after surgery, mice started a water restriction schedule to maintain 85–90% of free-drinking bodyweight for 5 days. The experimenter petted the mice 5 minutes per day for 3 days in a row and then started task training. All behavioral tasks were conducted during the dark cycle of mice.

The foraging task shuttle box has two chambers (10×10×15 cm) connected by a narrow corridor (45×5×15 cm, Fig 1a). A water port (1.2 mm O.D. steel tube, 3 cm above the floor) is attached to the end of one chamber defined as the reward zone, the other as the waiting zone. The position of the mouse in the shuttle box is tracked online with a custom MATLAB (2016b, MathWorks) program through an overhead camera (XiangHaoDa, XHD-890B). The experimental procedure control and behavioral event acquisition are implemented with a custom MATLAB program and an IC board (Arduino UNO R3).

### One-arm foraging task (pre-training)

A water-restricted mouse was put in the shuttle box for free exploration for up to 1 hour. When the animal started from the waiting zone through the corridor to the reward zone to lick the water port, 10 μl water was delivered by a step motor in 100 ms as a reward. A capacitor sensor monitors the timing and duration of licking. Then the animal returned to the waiting zone to re-initiate a new trial. Exiting from the waiting zone triggered an auditory cue (200 ms at 4 kHz sine wave with 90 dB) to signal the exit from the waiting zone. The time in the waiting zone was defined as the waiting duration. The training was conducted every day in a week. All mice learned to move quickly back and forth between two chambers to maximize the reward rate within one week.

### Delayed gratification task

From the second week, the volume of water reward is changed to a function proportional to the waiting time: 0∼2 s for 0 μl; 2∼4 s, 2 μl; 4∼6 s, 6 μl; 6∼8 s, 18 μl; >8 s, 30 μl as shown in Fig. 1A. There is no water delivered if the animal waits less than 2 sec. The training was conducted five days a week, from Monday to Friday.

### P_w_ Calculation

We divided all trials into two groups: Waiting Trials and Leaving Trials according to whether an animal to keep waiting or to leave in a given time duration such as 1 sec after each behavioral period. And then, we calculated the probability of waiting (P_w_) in this given time duration by the number of ‘Waiting Trials’ (*N*_*w*(*n*)_) and the number of ‘Leaving Trials’ (*N*_*L*(*n*)_) in the time window n:

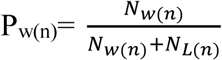

Then we can get the P_w_ for a given time duration:

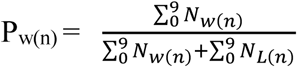

### Linear Mixed Model

We implemented the Linear Mixed Model Analysis using the open-source Python package “statsmodels” (https://www.statsmodels.org/stable/mixed_linear.html). The binary value of waiting or leaving during a specific behavioral period t_beh_ was set as the dependent factor (t_beh_=[1, 2), [2, 3), [3, 4), or [4, 5), unit: second); the fluctuation of momentary DA signal from its mean during a preceding period t_DA_ was set as a fixed effect (t_DA_=[0, 1), [1, 2), [2, 3), [3, 4), unit: second. Note that t_DA_ is always smaller than t_beh_); the animal identity and session numbers were set as a random effect (n=5 for each animal from the third week). The parameters of the model are estimated by restricted maximum likelihood estimation (REML).

### Optogenetic stimulation

Lasers, 473 nm for activation and 589 nm for inhibition, were coupled to the common end of a patchcord (200 µm O.D., 1-m long, 0.37 NA). The patchcord split through an integrated rotatory joint into two ends connecting to chronically implanted optical fibers (200 µm O.D., 0.37 NA) for bilateral light delivery. First, the mice were trained for 3 weeks to learn the delayed gratification task. Optical stimulation was delivered pseudo-randomly in ∼20% of behavioral trials in the test experiment. 20 ms square pulses at 10 Hz for activation or a continuous stimulation for inhibition were delivered. The laser was set to ON when the animal entered the reward zone and to OFF on the exit. The maximal laser stimulation was no longer than 16 seconds, even in the case a mouse stayed in the waiting zone longer than this time. Continuous laser power at the tip of splitting patchcord was about 10 mW for 473 nm laser and 8 mW for 589 nm laser, respectively.

### Random place performance test (RPPT)

After finishing optogenetic tests for delayed gratification, all mice took an RPPT. RPPT is carried on in a rectangular apparatus consisted of two chambers (30×30×30 cm) separated by an acrylic board. With an 8 cm wide door open, the mice could move freely between the two chambers. Before testing, each mouse was placed into the apparatus for 5-min free exploration. RPPT consists of two rounds of 10-min tests. First, we randomly assigned one chamber as a test chamber. Laser pulses were delivered in 20% possibility while the mouse entered the test chamber. The delivery of light, no longer than 16 sec, stopped while the mouse exited the test chamber. Next, we switched the chamber to deliver laser pulses. The laser output power and pulse length were set the same as optogenetic manipulations in the delayed gratification task.

### Fiber photometry recording

During the behavioral task training and test, we recorded the fluorescence signal of VTA dopaminergic (DAergic) neurons. The signal was acquired with a fiber photometry system equipped with a 488 nm excitation laser and a 505∼544 nm emission filter. The GCaMP6m signal was focused on a photomultiplier tube (R3896 & C7319, Hamamatsu) and then digitalized at 1 kHz and recorded with a 1401 digitizer and Spike2 software (CED, Cambridge, UK). An optical fiber (200μm O.D., 0.37 NA, 1.5-m long, Thorlabs) was used to transfer the excitation and emission light between recording and brain tissue. The laser power output at the fiber tip was adjusted to 5∼10 μW to minimize bleaching.

All data were analyzed with custom programs written in MATLAB (MathWorks). First, we sorted the continuously recorded data by behavioral trials. For each trial, the data spanned the range between 1 s before the waiting onset and 2 s after the reward. Before hooking the fiber to the mouse, we recorded 20 s of data and averaged as F_b_ as the ground baseline. For each trial, we averaged 1-sec data before the waiting onset as baseline F_0_ and then calculated its calcium transient as:

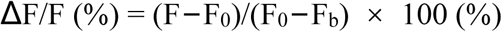

In the correlation analysis between VTA DAergic activity before waiting and waiting duration of mice, we used averaged 1-sec data before the waiting onset as the DAergic activity before waiting.

In the analysis of high-ramping and low-ramping DAergic activity, we compared the whole calcium signal of every trial with the average curve (the same length as the analyzed calcium signal) of all trials from one mouse in a single training day with paired t-test.

To facilitate presenting the data, we divided each trial data into four segments, including 1 s before waiting onset, waiting, running, and 2 s after rewarding. For comparing the rising trends, we resampled the data segments at 100, 100, 50, and 100 data points, respectively. In the delayed gratification task, the trial data were aligned to the waiting onset and presented by the mean plots with a shadow area indicating SEM of fluctuations.

### Reinforcement learning model

We investigate two potential scenarios. One was that the mouse decided on a waiting duration before entering the waiting area, and then waits according to the decided goal. The other scenario was that the mouse entered the waiting zone, and determined whether to wait or leave as an ongoing process throughout the whole waiting period. We called these two scenarios “*Decision Ahead”* and “*Continuous Deliberation”*, respectively, and formulated corresponding reinforcement learning based models for simulation using Python (Python Software Foundation, version 2.7. Available at https://www.python.org/).

#### Decision Ahead

Inspired by animal behavior, we simply set three optional “actions” with different expected waiting durations that could empirically cover the main range of animal’s waiting duration across training (*T*_a1_ = 1.65 sec for action1, *T*_a2_ = 2.72 sec for action2, *T*_a3_ = 4.48 sec for action3). These waiting durations were equally spaced on the log-time axis, consistent with Weber’s law (that is, ln(*T*_a1_) = 0.5, ln(*T*_a2_) = 1, ln(*T*_a3_) = 1.5). During the execution of action *a*_*i*_, we imposed additional noise to the timing so that the actual waiting time *τ*_ai_ for action *a*_*i*_ follows a Gaussian distribution on the log-time axis centered at the *T*_ai_, In ∼𝒩(In(*T*_ai_), 0.4^2^), *i* = 1, 2, 3. These settings allowed us to best capture the animal’s waiting performance in the model. For each trial, the agent chose action randomly based on three action values and a Boltzmann distribution (Softmax):

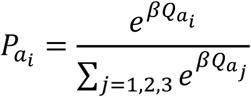

Where 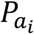 was the probability of choosing action *a*_*i*_ and waiting for *τ*_ai_. 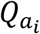 was the value for *a*_*i*_. *β* was the inverse temperature constant tuned to 5 according to our experimental data. After waiting, the agent would get a reward according to the same reward schedule used in our experiment. Each action value was updated separately during the reward delivery:

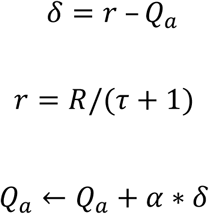

Where the reward prediction error *δ* was calculated by the difference between the hyperbolically discounted reward *r* (or “reward rate”, given by the absolute reward R dividing total time *τ*+1 for obtaining the reward, where *τ* was the waiting duration and the additional 1sec was the estimated delay for running between two zones) and the chosen action value *Q*_*a*_. The reward prediction error was then used to update the value of the chosen action. We tuned the learning rate α to 0.002 to fit the animal behavioral data.

#### Continuous Deliberation

In each trial, the agent would go through a series of hidden states, each lasting for 0∼2sec randomly according to a Gaussian distribution (mean at 1 sec). At each hidden state, the agent had two action options, either to keep waiting or to leave. If it chose to keep waiting, the agent would transition to the next hidden state, with the past time of the previous state cumulated to the whole waiting duration. If the choice was to leave, the cumulation would cease and a virtual reward dependent on the duration will be delivered, and then a new trial would begin from the initial state. The reward schedule was identical to that used for the animals during the experiments.

The action choice for the future was determined randomly by a Boltzmann distribution (SoftMax) and action values:

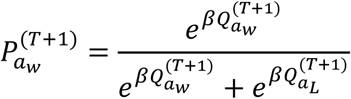

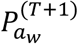 was the probability of choosing to wait for the next state *T* +1. 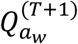 and 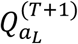 were the value of waiting and leaving, respectively, for state *T* + 1. *β* was the inverse temperature constant tuned to 5.

The action values for each hidden state *T* were updated by temporal difference learning algorithm (SARSA):

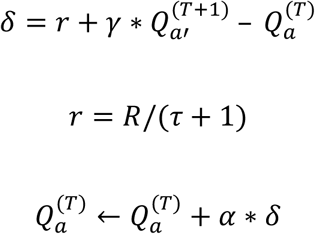

Where the future action *a*′ was determined by the Boltzmann distribution in the previous step. The current action *a* and the future action *a*′ could both be either waiting or leaving. The prediction error *δ* was calculated by the sum of reward rate *r* (r remained zero until the reward R was delivered. *τ*+1 was the total time for obtaining the reward, where *τ* was the waiting duration and the additional 1sec was the estimated delay for running between two zones) and the future action value 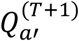 discounted by γ(γ = 0.9), minus the current action value 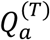. When *a* was leaving, the future action value 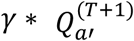would always be zero. This error signal *δ* was used to update 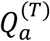 with learning rate α = 0.001.

As a Markov process, each state would be identical to the agent no matter how the state was reached or what the following actions are. So, we extracted the learned value of waiting as a time series along all the hidden states to compare with the averaged curve of VTA DAergic activity. For each trial, we also extracted the time series of the transient waiting value for a trial-wise analysis. Apart from the value of waiting, we could also extract the time series of RPE for each trial.

For optogenetics manipulation, we simulated it in the model after normal training was accomplished as in the animal experiments.

#### Value manipulation

In 20% trials of the stimulation session, the future waiting value throughout the whole waiting period was manipulated. The optogenetics activation was simulated as an extra positive value added onto the future waiting value, and the optogenetics inhibition corresponded to a proportional decrease of the future waiting value as follows:

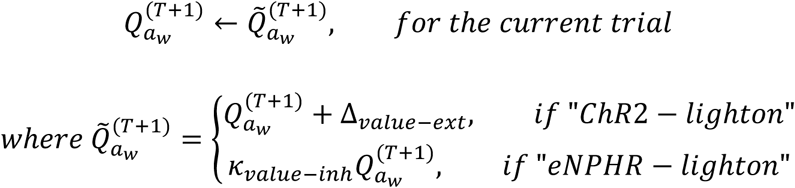

and, 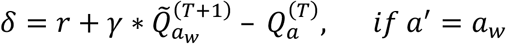

Here we set Δ_*value*−*ext*_= 0.15, and *K*_*value* −*inh*_ = 0.9, so that the change in averaged waiting duration in the simulated “light-on” trials can capture the magnitude of the instantaneous effect of optogenetic stimulations on the current trials. Using these parameters “calibrated” by the current trial effect, we were able to compare the stimulation effect on the light-off or the following trials in both real and simulated situations. Also note that if the future action was chosen as waiting, the manipulated value of waiting would be used in the RPE calculation and thus current action value updating as well.

#### RPE manipulation

Under this situation, in 20% trials of the stimulation session, instead of future waiting value, RPE (*δ*) was manipulated throughout the whole waiting period as follows:

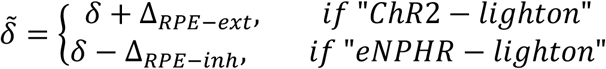

and, 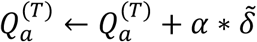
we set Δ_*PRE*−*ext*_= 15, and Δ_*PRE*−*inh*_= 20, which was calibrated by the current trial effect of real light stimulation.

To simulate the fluctuation in real DAergic signal, we simply multiplied the future waiting value during each state by a factor *σ*∼𝒩(1, 0.3^2^) (determined by the averaged signal-dependent noise magnitude / relative standard deviation for all momentary DAergic amplitudes), additionally to the original model (this is only implemented for figs. S8E∼F).

### Electrophysiological recordings

Adult (8-10 weeks) DAT-IRES-Cre knock-in male mice 4 weeks after injection with AAV2/9-EF1a-DIO-ChR2(H134R)-mCherry or AAV-DIO-eNpHR3.0-mCherry were anesthetized with an intraperitoneal injection of pentobarbital (100 mg kg^-1^) and then perfused transcardially with ice-cold oxygenated (95% O_2_**/**5% CO_2_) NMDG ACSF solution (93 mM NMDG, 93 mM HCl, 2.5 mM KCl, 1.25 mM NaH_2_PO_4_, 10 mM MgSO_4_·7H_2_O, 30 mM NaHCO_3_, 25 mM glucose, 20 mM HEPES, 5 mM sodium ascorbate, 3 mM sodium pyruvate, and 2 mM thiourea, pH 7.4, 295-305 mOsm). After perfusion, the brain was rapidly dissected out and immediately transferred into an ice-cold oxygenated NMDG ACSF solution.

Then the brain tissue was sectioned into slices horizontally at 280 mm in the same buffer with a vibratome (VT-1200 S, Leica). The brain slices containing the VTA were incubated in oxygenated NMDG ACSF at 32° for 10∼15 min, then transferred to a normal oxygenated solution of ACSF (126 mM NaCl, 2.5 mM KCl, 1.25 mM NaH_2_PO_4_, 2 mM MgSO_4_·7H_2_O, 10 mM Glucose, 26 mM NaHCO_3_, 2 mM CaCl_2_) at room temperature for 1h. A slice was then transferred to the recording chamber, which was submerged and superfused with ACSF at a rate of 3 ml/min at 28°. Cells were visualized using infrared DIC and fluorescence microscopy (BX51, Olympus). VTA DAergic neurons were identified by their fluorescence and other electrophysiological characteristics. Whole-cell current-clamp recordings of VTA DAergic neurons were made using a MultiClamp 700B amplifier and Digidata 1440A interface (Molecular Devices). Patch electrodes (3-5 MΩ) were backfilled with internal solution containing (in mM): 130 K-gluconate, 8 NaCl, 10 HEPES, 1 EGTA, 2 Mg·ATP and 0.2 Na_3_·GTP (pH:7.2, 280 mOsm). Series resistance was monitored throughout the experiments. For optogenetic activation, blue light was delivered onto the slice through a 200-μm optical fiber attached to a 470 nm LED light source (Thorlabs, USA). The functional potency of the ChR2-expressing virus was validated by measuring the number of action potentials elicited in VTA DAergic neurons using blue light stimulation (20 ms, 10 Hz, 2.7 mW) in VTA slices. For optogenetic inhibition, yellow light (0.7 mW) was generated by a 590 nm LED light source (Thorlabs, USA) and delivered to VTA DAergic neurons expressing eNpHR3.0 through a 200-μm optical fiber. To assure eNpHR-induced neuronal inhibition,whole-cell recordings were carried out in current-clamp mode and spikes were induced by current injection (200 pA) with the presence of yellow light. Data were filtered at 2 kHz, digitized at 10 kHz, and acquired using pClamp10 software (Molecular Devices).

### Immunostaining

Mice were deeply anesthetized with pentobarbital (100 mg/kg, i.p.), following saline perfusion through the heart. After blood was drained out, 4% paraformaldehyde (PFA) was used for fixation. Then the head was cut off and soaked in 4% PFA at room temperature overnight. The brain was harvested the next day, post-fixed overnight in 4% PFA at 4°C, and transferred to 30% sucrose in 0.1 M PBS, pH 7.4 for 24∼48 h. Coronal sections (20 µm) containing the VTA were cut on a cryostat (Leica CM3050 S). The slides were washed with 0.1 M PBS, pH 7.4, incubated in blocking buffer (0.3% Triton X-100, 5% bovine serum albumin in 0.1 M PBS, pH 7.4) for an hour, and then transferred into the primary antibody (rabbit anti-tyrosine hydroxylase antibody, 1:1,000; Invitrogen) in blocking buffer overnight at 4°C. The sections were washed three times in 0.1 M PBS, then incubated with donkey anti-rabbit IgG H&L secondary antibody (conjugated to fluor-488 or fluor-594, 1:1,000; Jackson ImmunoResearch) at room temperature for 2 h. The nucleus was stained with DAPI (4’,6-diamidine-2-phenylindole). Sections were mounted in glycerine and covered with coverslips sealed in place. Fluorescent images were collected using a Zeiss confocal microscope (LSM 880).

### Quantification and statistics

All statistics were performed by MATLAB (R2016b, MathWorks) and Python (V2.7, Python Software Foundation) routines. Data were judged to be statistically significant, while the P-value less than 0.05. Asterisks denote statistical significance *p < 0.05; **p < 0.01; ***p < 0.001. Unless stated otherwise, values were presented as Mean ± s.e.m..

## Acknowledgments

We thank Drs. M. Lou, W. Ge, Y. Rao, W Zhou, and B Min for comments on the manuscript. This work was supported by the National Natural Science Foundation of China (grant nos. 31922029, 31671086, 61890951, and 61890950 to J.H.), a Shanghai Pujiang Talent Award (grant no. 2018×0302-101-01 to W.S.).

## Author contributions

W.S. and J. H. oversaw the whole project. W.S., J. H. & Z. G. designed the experiments. Z. G. & C. L. performed all animal experiments. Z. G. & H. W. analyzed the data. H. W. & S. F. performed the computational modeling under the supervision of X. J. W.. M. C. performed the electrophysiological recordings. W.S., J. H., H. W. & Z. G. wrote the paper with the participation of all other authors.

---

**Fig. S1.**
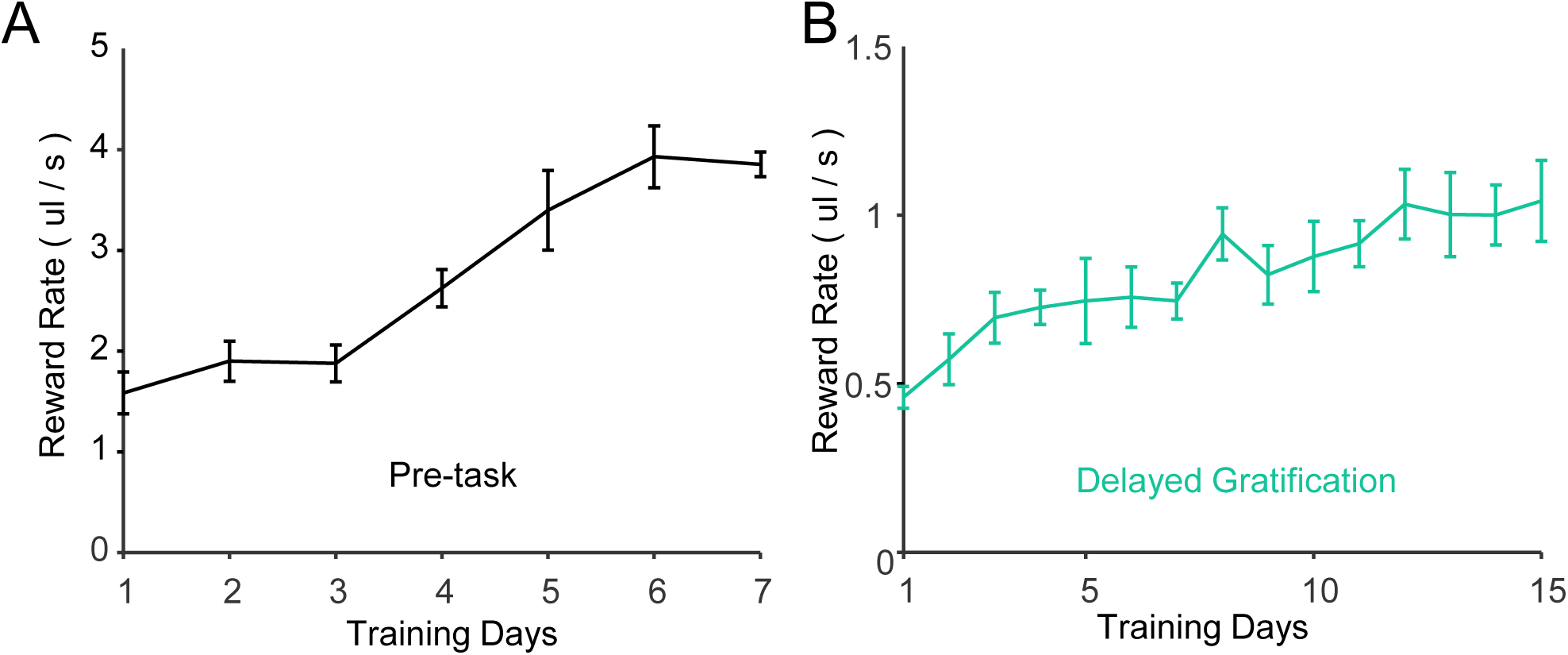
The reward rate during behavioral training. (**A**-**B**) The reward rate both increased in pre-task training (**A**) and delayed gratification training (**B**). All error bars represent the s.e.m..

**Fig. S2.**
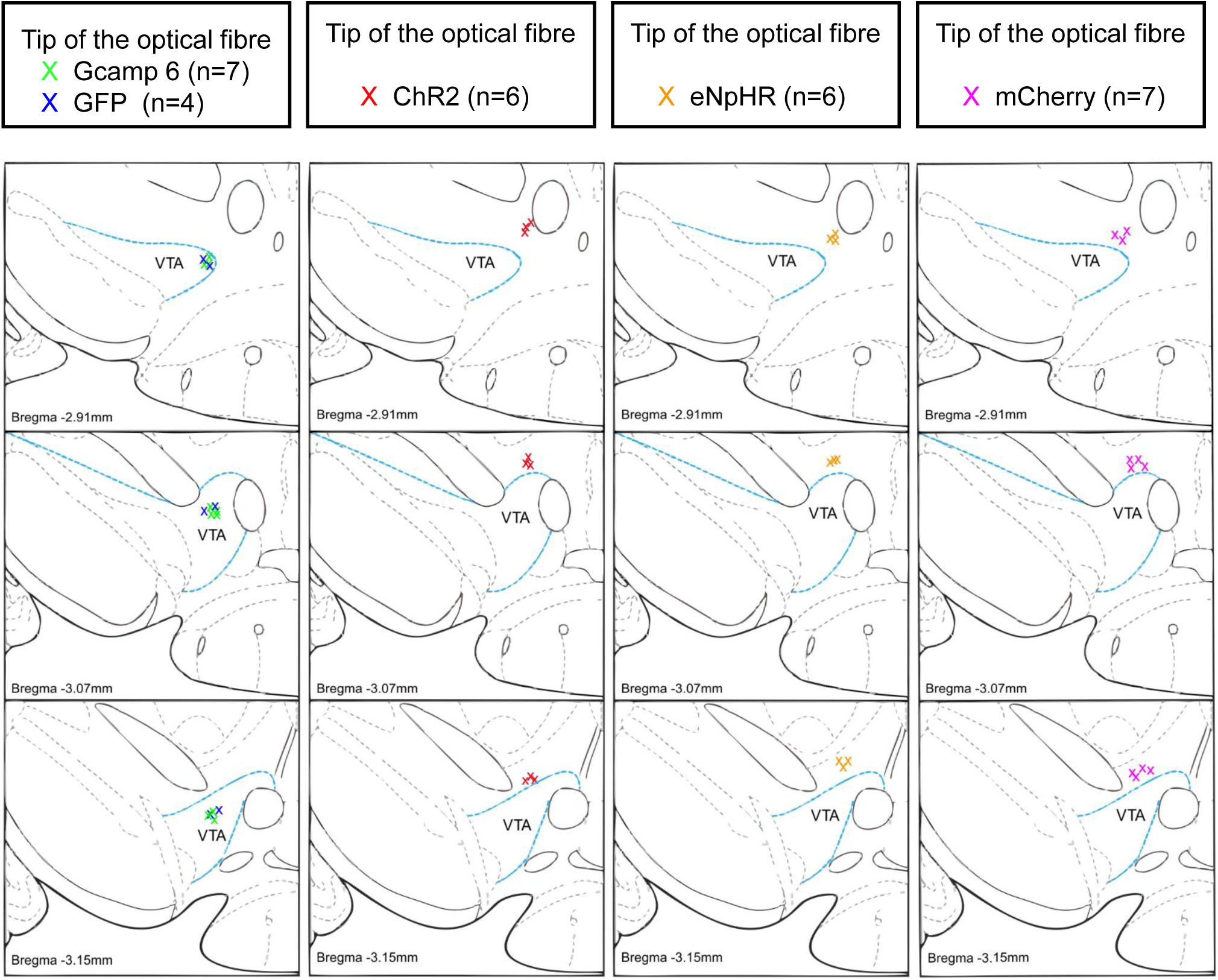
Tip Positions of optical fibre in GCaMP6m (green, n = 7), GFP (blue, n = 4), ChR2 (red, n = 6), ENpHR (yellow, n = 6) and mCherry (magenta, n = 7) shown as coordinates in the mouse brain atlas.

**Fig. S3.**
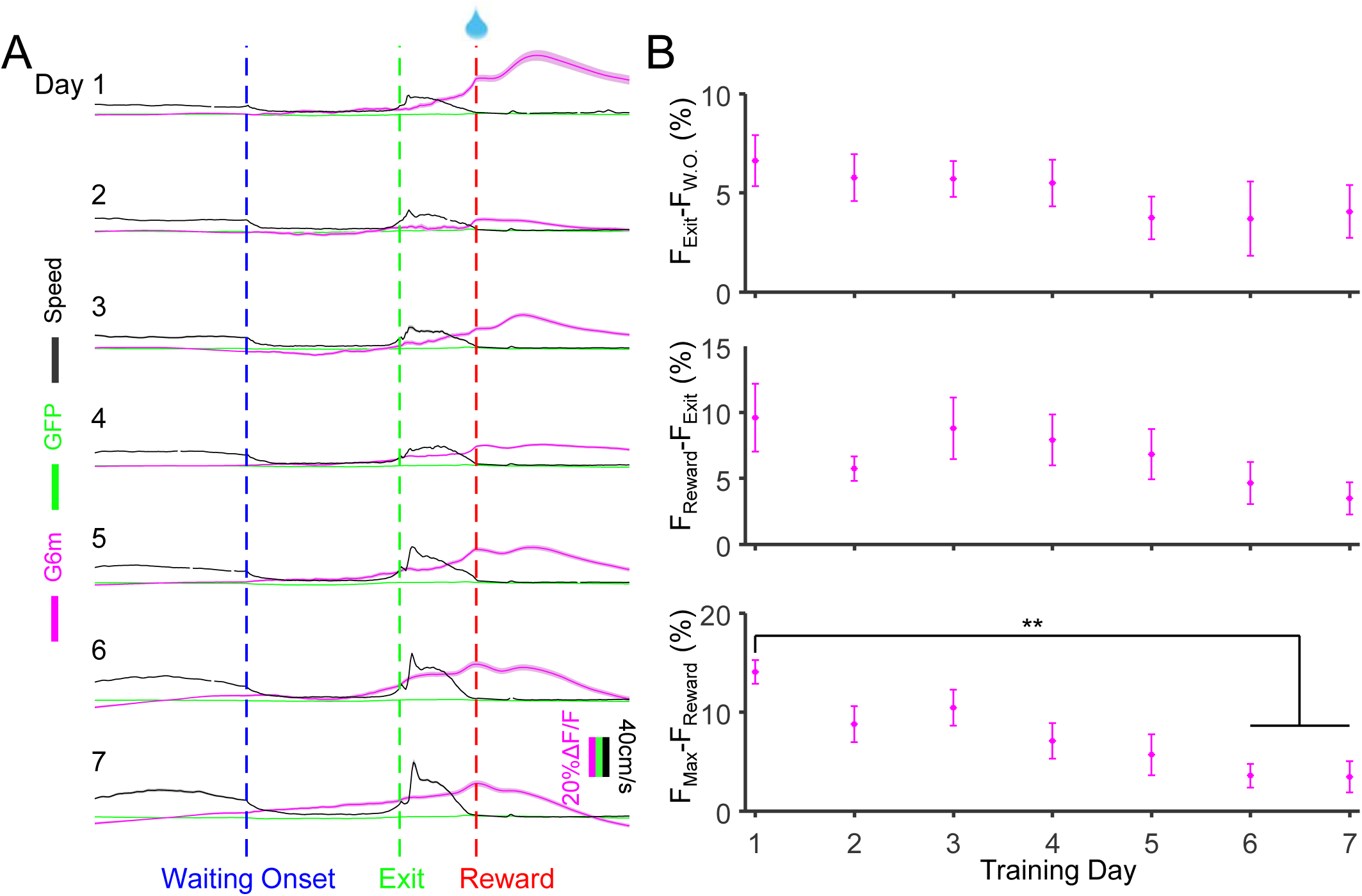
The calcium and GFP signals of VTA DA neurons in behavioral tasks. (**A**) The GFP (green), calcium signals (magenta) of VTA DA neurons, and mouse running speed (black) in 7 days of pre-training. (**B**) The calcium signals of VTA DA neurons were changing with pre-training progress. The max calcium signals after the mouse received water rewards significantly decreased in day 6&7 (p<0.01, Friedman test). All error bars represent the s.e.m..

**Fig. S4.**
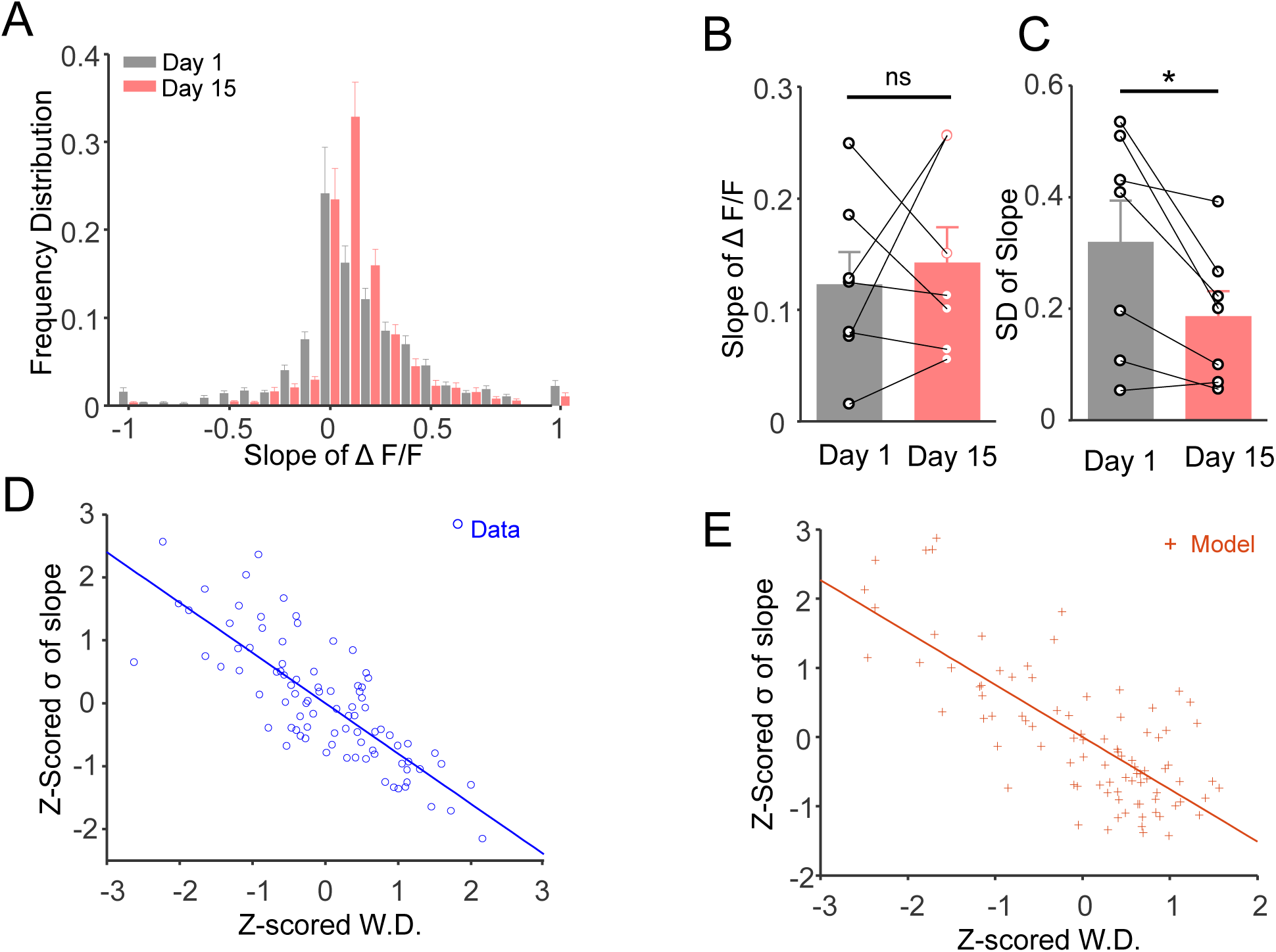
The ramping dopamine activity became more stable with delayed gratification training. (**A**) The Frequency distribution of the slope of Z-scored ΔF/F during waiting (gray: day 1, red: day 15, n = 7). (**B**) The slope of ΔF/F had no difference between day 1 and day 15 (p=0.63, paired Student’s t-test). (**C**) The standard deviation(σ) of the slope of ΔF/F in day 15 (0.32±0.07) was significantly decreased than that in day 1 (0.17±0.05, p=0.03, paired Student’s t-test). (**D**-**E**) The z-scored σ of ΔF/F and z-scored waiting durations were negatively correlated both in the experimental data (d, blue, r = −0.80, p<0.001) and RL model (e, red, r = −0.76, p<0.001). W.D. is for waiting duration. All error bars represent the s.e.m..

**Fig. S5.**
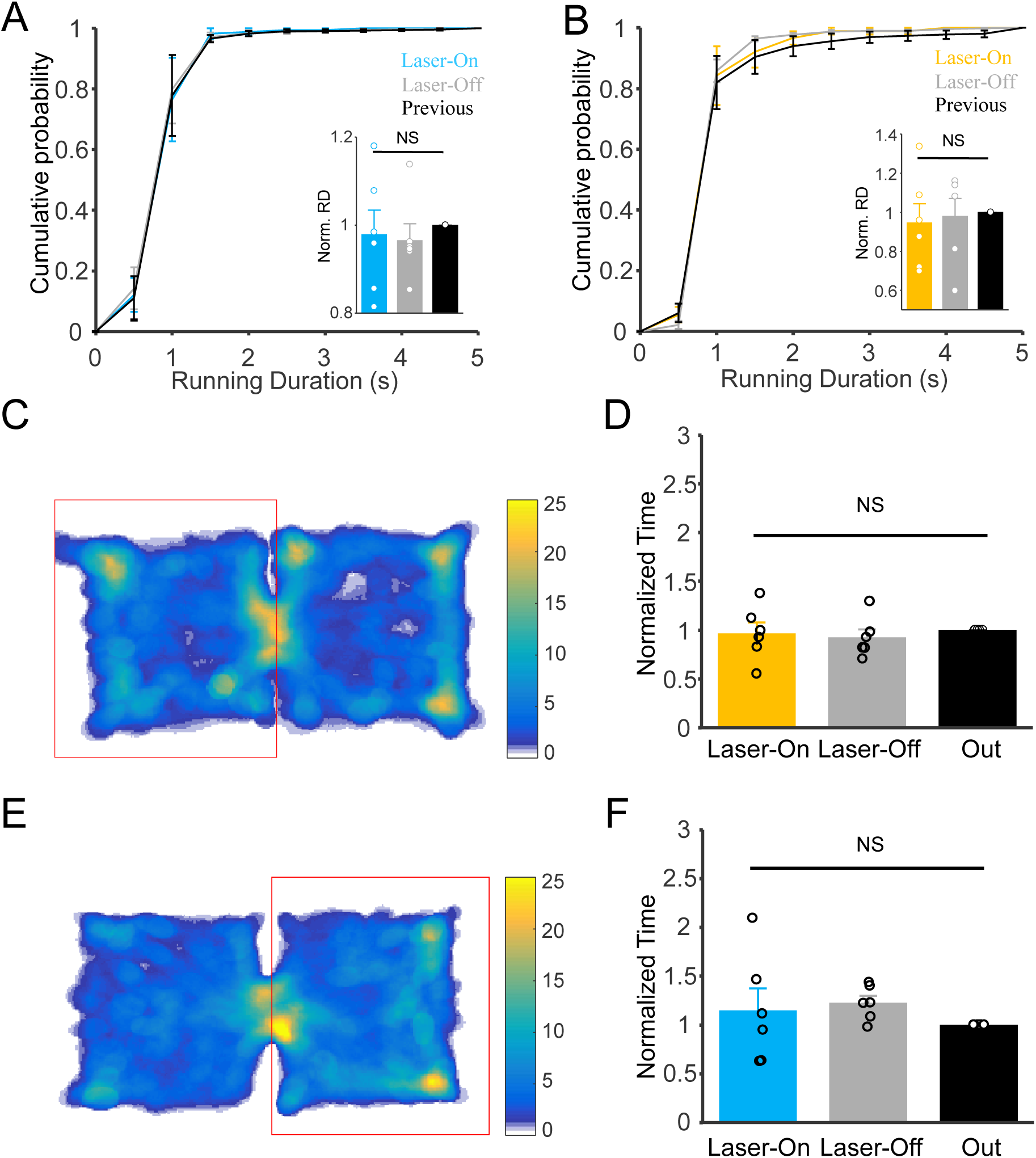
Optical activation or inhibition of VTA DA activity didn’t change the motivation of mouse in the delayed gratification tasks and the duration the mouse stayed in the given box in RPPT. (**A**) The running duration didn’t change while the VTA DA neurons were optically activated (F=0.20, p=0.82, one-way ANOVA). (**B**) The result was the same as Figure A while optical inhibiting the VTA DA neurons (F=0.12, p=0.88, one-way ANOVA). (**C**) The heatmap of mouse traces in RPPT in which the VTA DA neurons were optically activated pseudo-randomly in 20% probability while the mouse entered into the given box (red rectangle). (**D**) The Z-scored duration that the mouse stayed in the given box while the VTA DA neurons were activated (Laser-On In) had no significant difference (F=0.75, p=0.44, one-way ANOVA, n=6) with the un-inhibited durations (Laser-Off In) and durations in another box (Out). (**E**) The heatmap of mouse traces as shown in Figure C while inhibiting the VTA DA neurons in the given box (red rectangle). (**F**) Optical inhibiting the VTA DA neurons also didn’t change the duration the mouse stayed in the given box (F = 0.17, p = 0.73, one-way ANOVA, n=6). All error bars represent the s.e.m..

**Fig. S6.**
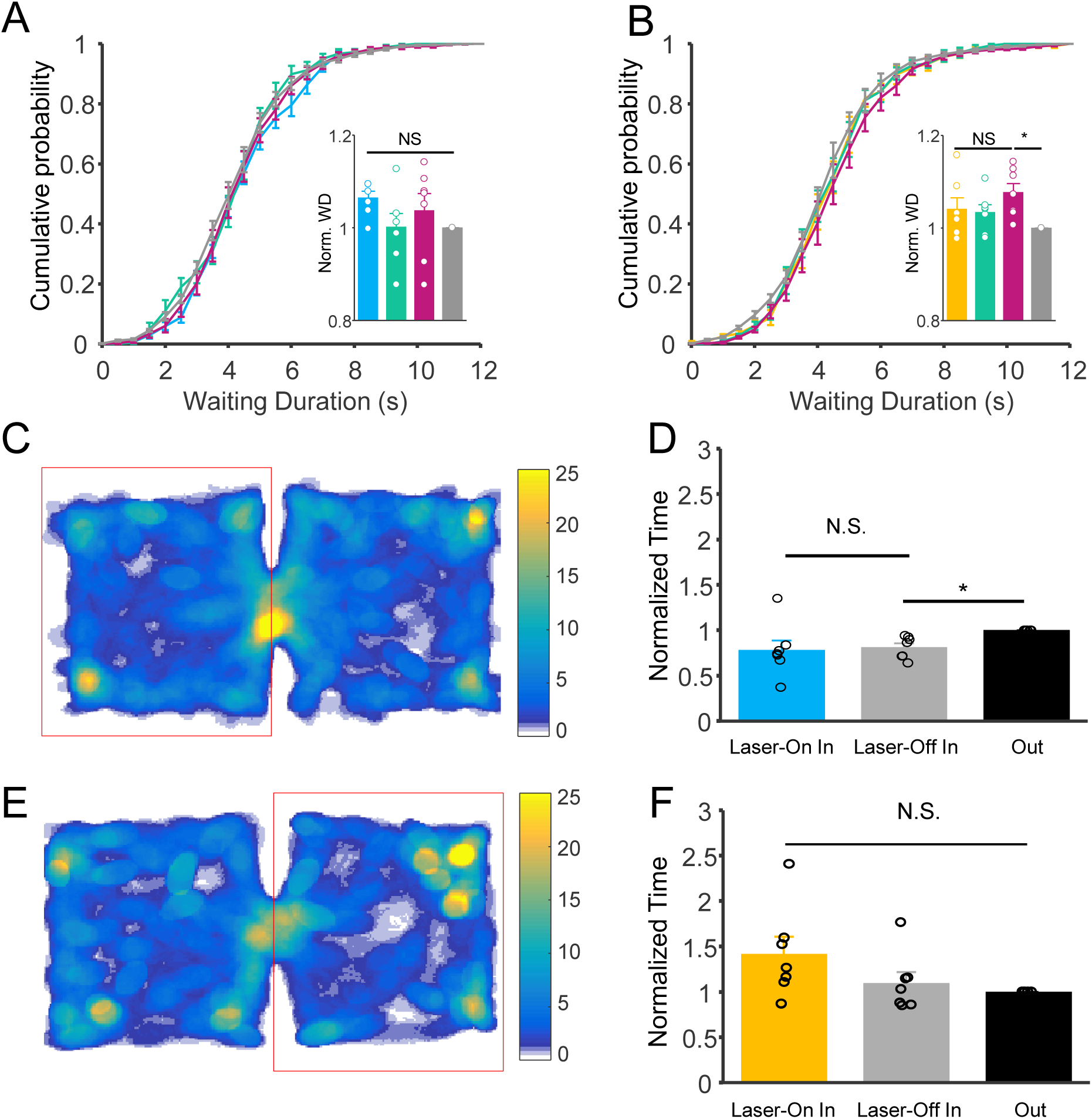
Optogenetic manipulation of DAT-Cre mouse expressed mCherry in the delayed gratification tasks and RPPT. (**A**) Waiting durations in 473nm laser delivered trials (blue) are not different compared with those of all other trials (p=0.17, Friedman test, n=7). (**B**) Waiting durations of 589nm laser un-delivered trials (magenta) slightly increased compared with the waiting duration of the previous day (p=0.02, Friedman test, n=7). (**C**) Heat-map of mouse traces in RPPT in which the VTA of the mouse was delivered 473nm laser pseudo-randomly in 20% probability while the mouse entered into a randomly chosen box (red rectangle). (**D**) Mean durations that the mouse stayed in chosen box while the laser delivered (Laser-On In), laser off (Laser-Off In) and the other box. There is no significant difference in waiting duration between Laser-On In and Laser-Off In (F=3.54, p=0.09, one-way ANOVA, n=7). (**E**) Heatmap of mouse traces same as shown in (c) while the mouse was delivered 589nm laser in a randomly chosen box (red rectangle). (**F**) Mean durations that the mouse stayed in chosen box while 589nm laser delivered (Laser-On In), laser off (Laser-Off In), and in the other box. 589nm laser delivering to mCherry mouse didn’t alter waiting duration mouse stayed in any boxes under all experimental conditions (F=2.64, p=0.14, one-way ANOVA, n=7). All error bars represent the s.e.m..

**Fig. S7.**
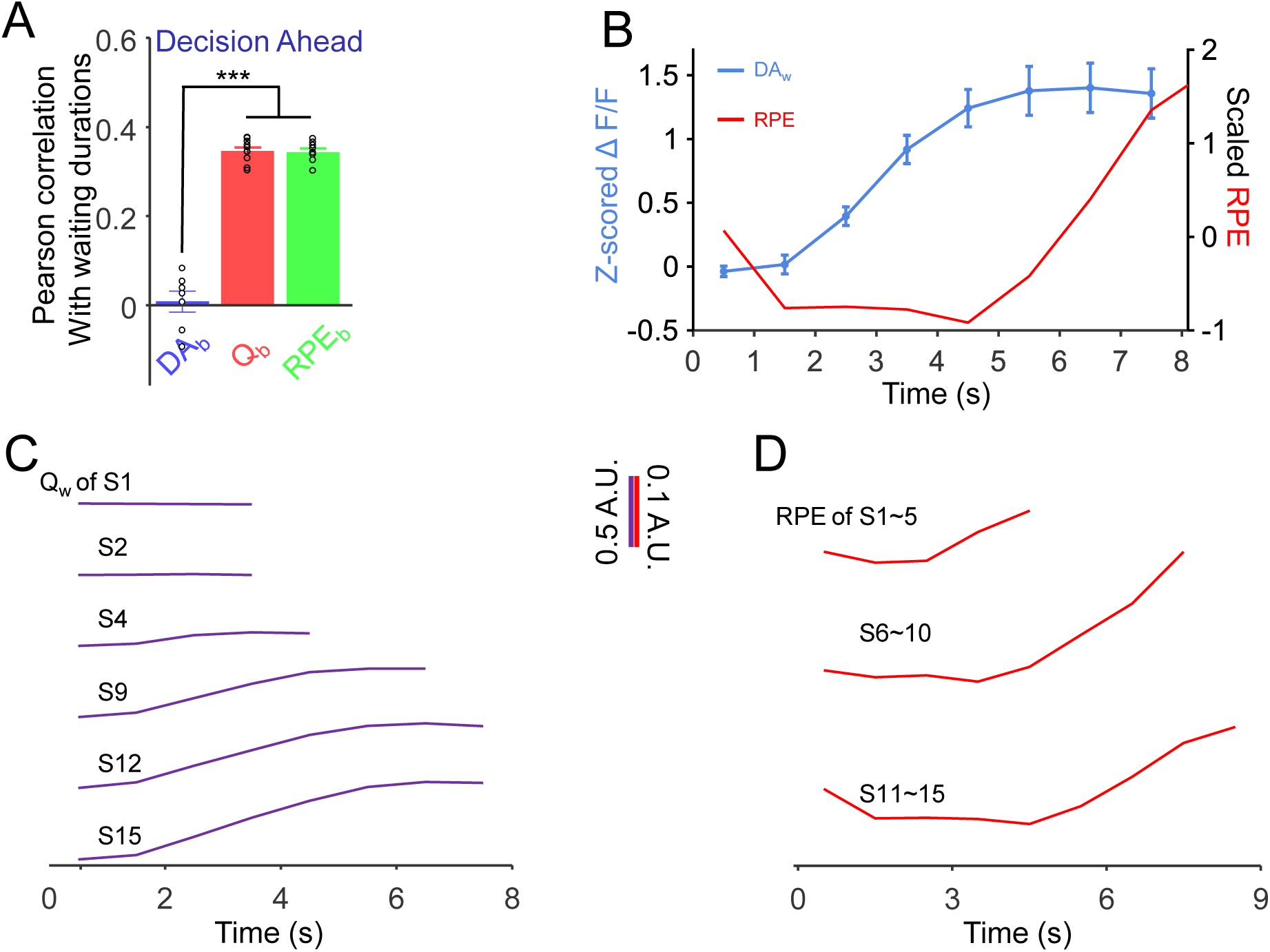
The Data from RL models. (**A**) The correlation coefficient of mean DA activity 1 s before waiting (DA_b_), the value of action (Q_b_), and RPE of action (RPE_b_) before waiting in the Decision Ahead model, with waiting durations. The correlation of V_b_ (r = 0.35±0.01, p=0.001, n=10, Pearson correlation) and RPE_b_ (r = 0.34±0.01, p=0.001, n=10, Pearson correlation) with waiting duration were significantly (p<0.001, Kruskal-Wallish test) higher than the CC of DA_b_ (r=0.01±0.02, p=0.36, n=7) with waiting durations. (**B**) Plots of Z-scored ΔF/F values (DA_w_, light blue) at 0.5s before the waiting ended and RPE of waiting (RPE_w_). There were no significant correlation bewteen DA_w_ and RPE_w_ (r = 0.34, p = 0.41, Pearson correlation). (**C**-**D**) Value of waiting (Q_w_) (**C**) and RPE (**D**) changed in the Continuous Deliberation model. All error bars represent the s.e.m..

**Fig. S8.**
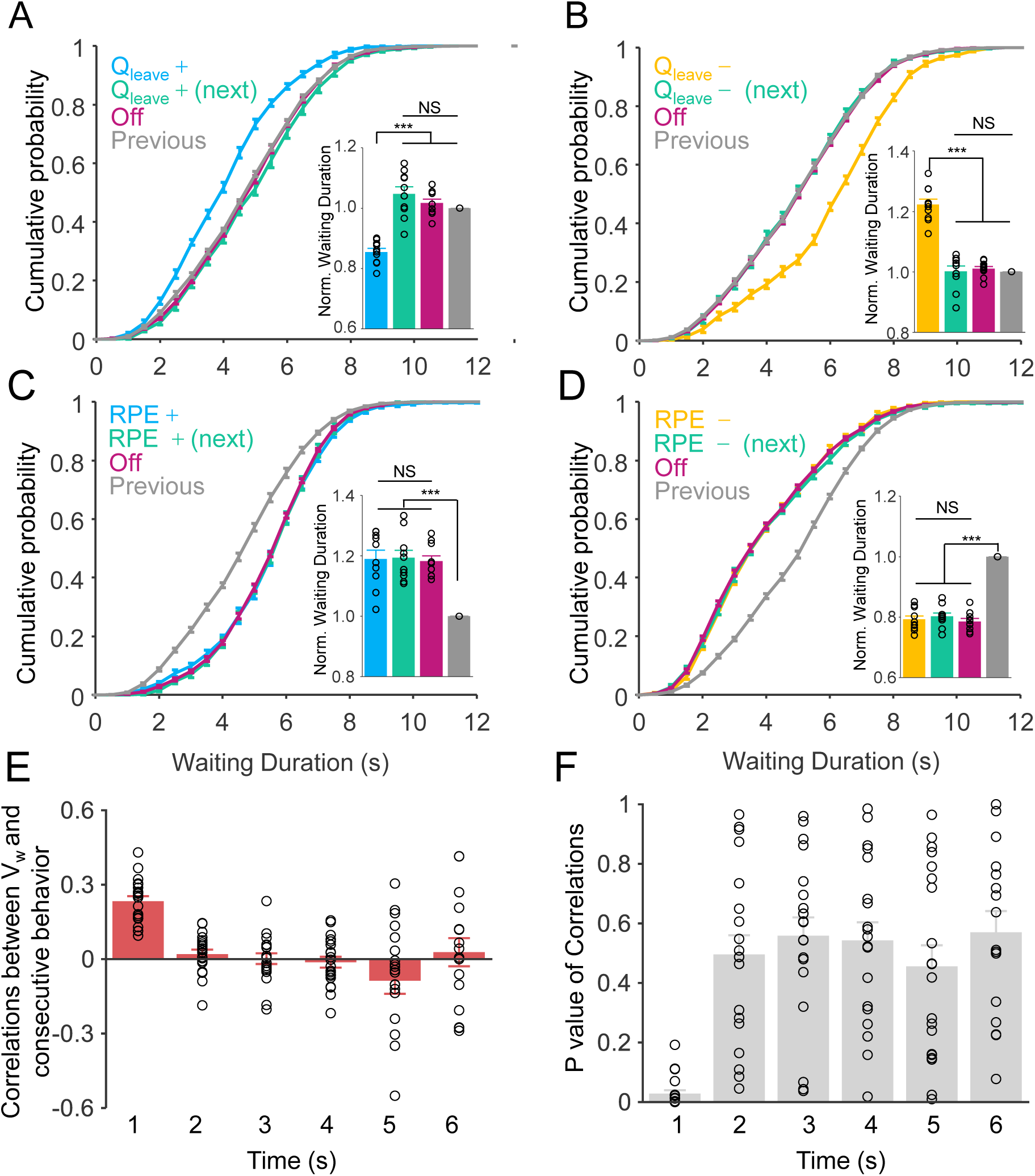
Manipulation of value of leaving and RPE in RL model. (**A**-**B**) Either increasing or decreasing the value of leaving (Q_leave_) in the continuous deliberation model as with the manipulation of Q_wait_ induced the opposite results compared with the optogenetics manipulating DAergic activity (increasing Q_leave_ in **A**: p<0.001, Friedman test, n=10; decreasing Q_leave_ in **B**: p <0.001, Friedman test, n=10) and had no influences on other trials (**A-B**, p>0.999, Friedman test, n=10). (**C**-**D**) Either increasing (**C**) or decreasing (**D**) the RPE in the continuous deliberation model as with the experimental data, alters the waiting durations in the same direction in all trials, whether or not the RPEs-manipulation (increasing RPEs in c: p<0.001, Friedman test, n=10; decreasing RPE in (**D**): p <0.001, Friedman test, n=10). (**E**) The value of waiting is only positively correlated (0.23±0.02, p =0.03±0.01, n =20) with the adjacent behavior in the *Continuous Deliberation* RL model. (**F**) The p values of correlation coefficients in **E**. All error bars represent the s.e.m..

